# Analysis of Age-Specific Dysregulation of miRNAs in Lung Cancer Via Machine learning: Biomarker Identification and Therapeutic Implications in Patients Aged 60 and Above

**DOI:** 10.64898/2026.02.12.705605

**Authors:** Ahmed Hasan, Arfa Muzaffar

## Abstract

Lung cancer is the leading cause of cancer-related mortality worldwide, predominantly affects older individuals, with **non-small cell lung cancer (NSCLC)** comprising 85% of cases. Despite advancements in diagnosis and treatment, prognosis for elderly patients remains poor. This study investigates the role of microRNAs (miRNAs) involved in lung cancer, focusing on individuals aged 60 and above. RNA sequencing data from **The Cancer Genome Atlas (TCGA)** was used to conduct differential expression analysis of miRNA profiles from elderly and senile patient groups. Results showed that out of 1,881 miRNA profiles, 801 were found to be differentially expressed. Filtering for significance identified that 25 miRNAs, with *hsa-mir-1911* upregulated and 24, including *hsa-mir-196a* and *hsa-mir-323b* found to be downregulated. Studies showed that these miRNAs play roles in apoptosis, senescence, and inflammation. Another Experimental approach in this study, used Machine learning analysis which highlighted key miRNAs, including *hsa-mir-181b*, *hsa-mir-542*, *hsa-mir-450b*, *hsa-mir-584*, and *hsa-mir-21* as crucial in lung cancer biology. Moreover, Functional enrichment analysis revealed their involvement in gene silencing, translational repression, and RNA-induced silencing complex (RISC) regulation. This research identifies the association of miRNAs and aging in lung cancer and finds potential biomarkers that can be helpful in early diagnosis and targets for personalized therapies.

## INTRODUCTION

lung cancer is currently the most prevalent cancer globally and is responsible for deaths of around 1.8 million people each year (Bray et al., 2020). Lung cancer can be categorized into: small cell lung cancer, and non-small cell lung cancer NSCLC which comprises 85% of the total cases of lung cancer: the common subtype is adenocarcinoma. (Siegel et al., 2022)

De novo molecular technologies have in recent years led to early detection of the disease and better treatment regimens but lung cancer still poses a bleak picture with a 5-year survival rate pegged at 20% (American Cancer Society, 2023). These figures provide further emphasis to the plight of diagnosing and effectively treating for better patient prognosis.

Lung cancer management is even more complex owing to factors such as age, which is a risk factor for lung cancer (Patz et al., 2016). Molecular changes in this group include age dependent genomic instability in oncogenes and tumor-suppressor genes, presence of comorbid illness, and advanced stage of tumor at the time of diagnosis (Yancik et al., 2005). This underlines the need to find molecular predictors that would shed more light on the different biology of lung cancer in the elderly and may direct the development of individualized therapies.

MicroRNAs (miRNAs) are a group of small, non-coding RNA molecules that can bind target messenger RNA (mRNA) at its 3’ untranslated region (UTR) thereby promoting mRNA degradation or inhibition of translation (Bartel, 2004). Elucidating the functions of these molecules has highly vital roles in several biological activities such as cell growth, development, death, and movement (He & Hannon, 2004). Aberrant expression of miRNAs has been reported to be involved in many human cancers, including lung cancer, and can act as oncogenes or as tumor suppressor genes (Esquela-Kerscher & Slack, 2006). That miRNAs are stable in biological fluids including blood and serum also support their possibilities as noninvasive biomarkers for cancer diagnosis, prognosis and treatment outcome (Mitchell et al., 2008).

Some prior research has focused on analyzing the potential of miRNAs in lung cancer. For example, miR-21 has been reported to be an oncomiR that enhances tumorigenesis through downregulation of tumor suppressor genes including PTEN and PDCD4 (Chan et al., 2005). Conversely, the let-7 family of miRNA functions as a tumor-suppressive agent by targeting for degradation or translational repression of oncogenes like RAS and HMGA2 (Johnson et al., 2005). However, specific knowledge regarding age-specific perturbation of miRNA and its role in lung cancer progression in elderly patients is scarce. This shortestcoming is of much concern given the fact that there has been a significant rise in incidence of lung cancer among the aging population (de Groot et al., 2018).

In this regard, the present study aims to make progress towards filling this gap by examining and describing at least 10 miRNAs that are differentially expressed in patients with lung cancer who are 60 years or older. Given the RNA-seq data obtained from TCGA, the differential expression analysis was conducted to recognize miRNAs which play a role in tumor progression in this age group. The prediction showed that the expression of the following miRNAs may be related with lung cancer, including hsa-let-7b, hsa-mir-196b, hsa-mir-381, and hsa-mir-484. The above miRNAs were selected because of their expression profile, functional importance, and role in cancer as depicted in the following literature.

### Aim and Objectives

#### Aim of the study

Although there are numerous preexisting works which explain certain miRNAs that function during the development of lung cancer, the correlation between the global dysregulation of miRNAs and their dysregulation based on age remains very limited in lung cancer patients, especially in the senior population. Normal aging is characterized by multiple molecular and cellular alterations such as enhanced oxidative stress, inflammation and altered epigenetic profile, all of which can impact the tumor microenvironment profoundly (López-Otín et al., 2013). These changes not only impact on the levels of gene expression but also the function of miRNAs and thus tumor phenotype and response to treatment could be different in older people. However, these changes have been well described in the literature and, surprisingly, most of the miRNA investigations on lung cancer fail to consider age factor while some address it mainly to young subjects thus; negating the results in reference to elderly population.

The second major limitation is the lack of a confluence of multiple bioinformatics analyses to align miRNA expression patterns and possible pathways implicated in older patients. Present day experiments utilize single model systems and techniques, thus obviating feedback regulatory systems as well as the pathway level information, which could offer a comprehensive understanding of miRNA roles. Besides, the clinical approach of defining miRNAs as biomarkers in elderly persons is still limited, which in turn hampers their usability from bench to clinic.

#### Objectives of the Study

The overall research question of this study can be stated as follows: What specific miRNA has an impact on lung cancer in patients more than sixty years old? Thus, based on the RNA-seq data analysis and a literature review of the existing literature, this work aims to fill important gaps in the comprehension of miRNA-associated regulation in older patients. Specifically, the objectives are as follows:

##### Identification of Dysregulated miRNAs

Compare miRNA activity in lung cancer and healthy lung samples to find regulatory changes specific to patients over 60 years old.Detects age-related expression changes in miRNAs between patients from two different age groups.

##### Pathway Analysis and Functional Insight

Show which key biological pathways changed when miRNAs do not work properly in older lung cancer patients.

Bioinformatics analysis helps track how miRNAs interact with genes and reveals which cancer-promoting and cancer-blocking signaling pathways they modify.

##### Biomarker Potential Evaluation

- Study the ability of miRNAs to correctly detect when tissues are cancerous or not.
- Measure miRNA expression levels to find connections between their patterns and how patients fare over time and how well they react to treatments.
- Find miRNAs that medical teams might target to create therapeutic strategies employing miRNA mimics or inhibitors in research.

### Significance of Age-Specific Research

Knowledge about lung cancer’s molecular structure in older patients helps build better tailored treatment approaches. The aging process impacts multiple cellular functions including the immune system’s ability to fight off disease plus helps damage build up inside cells while adjusting their energy usage styles. This study evaluates microRNA expressions to find diagnostic signs and treatment possibilities that directly benefit older patients who account for more lung cancer cases than studies typically focus on.

#### 1.2.1 Anticipated Impact

This study has the potential to bridge critical gaps in miRNA research by providing:

- We present an extensive list of age-correlated miRNAs that affect lung cancer.
- We discovered how aging affects biological processes in older patients which benefits both aging cancer patients and clinical research.
- Our research sets the groundwork for drug trials using specific microRNA treatments designed for older lung cancer patients

##### Methodological Approach

We examined RNA-seq samples from the TCGA-LUAD lung cancer patient cohort to/reach our findings. We limited the dataset to patients over 60 years old to study age-related patterns. Our differential expression analysis used the DESeq2 package from R (Love et al., 2014) to detect miRNA expression changes in the RNA-seq data. DAVID and KEGG databases helped us identify biological pathways connected to these miRNAs by following the approach of functional enrichment analysis as presented by (Huang et al. in 2009).

Our findings backed up previous research evidence to verify the identified miRNAs’ functions in lung cancer development. Researchers confirmed let-7b’s tumor-suppressing function by showing it reduces both lung cancer cell growth and spread in lab experiments (Liang et al., 2019). Results from Song et al., 2019 proved that miR-196b behaves as an oncogene by showing it contributes to poorer survival rates in NSCLC patients.

###### 1.4 Significance of the Study

We expand current studies about cancer miRNAs by studying how they function differently in lung cancer patients of varying ages. Research on let-7b miR-196b miR-381 and miR-484 miRNAs shows these molecules play key roles in the molecular changes that lead to lung cancer progression in aging patients. These findings show how miRNAs can serve as both treatment targets and disease markers to create personalized cancer therapies

## MATERIALS AND METHODS

### Data Downloading

TCGA-Biolinks was used to download miRNA Expression profiles and associated metadata of Lung Cancer patients from TCGA-LUAD Project from the GDC portal. (Colaprico et al., 2016).

### Filtering and Merging

Raw data consisted of 585 age samples associated with 1882 miRNA Expression profiles. The Elderly samples were 269 and the senile samples were 101. Upon merging, only 204 Elderly and 76 senile samples were left.

### Differential Expression Analysis

801 Differentially expressed miRNAs after the original data set of 1881 miRNAs using the Deseq2 package. The differential expression was run by grouping ages as previously defined during filtering of the data i.e. 60-75 and 75-90.

Results of Differential expression analysis were analyzed to identify 25 significant miRNAs by filtering based on padj value < 0.05 and log2 fold change >1.

### Machine Learning based Analysis

#### Data Acquisition and Preprocessing

The merged data was used to train the machine learning algorithm.R loaded the dataset through read.csv() while each sample row maintained a distinct identity and each expression level served as its own column. The dataset contained a final column which identified the age group categories.

We applied quantile normalization using normalize.quantiles() from preprocessCore to maintain consistent measurement results across different samples. Each sample’s statistical distribution becomes normalized through this method which eliminates technical variability biases.

#### Feature Selection

Feature selection improved both model efficiency while reducing system complexity by applying cross-sample variance computation on miRNA sequences. By using the apply() function researchers retrieved variance values before selecting the top 500 miRNAs for following investigation. We selected the most informative miRNAs through this step while filtering out variability-low features to eliminate potential noise factors.

#### Machine Learning Model

The Random Forest classification algorithm from the randomForest package enabled supervised machine learning implementation. The following steps were performed:

This data consisted of predictor top 500 miRNAs and response terms related to age classification.

The trained Random Forest model used randomForest() to build its structure with 500 decision trees (ntree = 500) at default split variable selection mode.

Recorded importance scores from the model enabled extraction through the importance() function while sorting miRNAs according to their contributions to prediction accuracy.

#### Model Evaluation

Predictions from the model ran against the training dataset through the predict() feature. The confusionMatrix() function from caret generated an assessment tool to analyze classification precision and recall alongside accuracy rates as well as general error estimation. A supplementary measure of model performance emerged through the usage of the out-of-bag (OOB) error estimate.

### Visualization and Interpretation of DEA And ML results

#### Visualization of ML results

Feature interpretation relied on Mean Decrease in Gini index values to identify the key miRNAs. A heatmap created with pheatmap showed how expression levels changed between different samples. The implementation of ggplot2 feature importance plots helped determine which miRNAs acted as the strongest predictors for age group classification.

The proposed research methodology uses logical stages of statistical preprocessing followed by feature selection along with machine learning and visualization to generate significant biological findings from miRNA expression data.

#### Visualization of DEA results

To elucidate the results:

- **MA Plot**: log2 fold changes were plotted against mean expression levels using MA Plot to highlight important identified miRNAs.
- **Volcano Plot**:Enhanced Volcano plot was used to show the distribution of all the miRNAs identified in the differentially expressed and segregating significant, Non-significant and NA micro RNAs using different colours respectively.

##### miRNA Target Prediction and Functional Annotation

The significant micro RNAs were then mapped to their genes using the biomart package and Ensembl’s Database.

###### Gene Ontology Enrichment Analysis

To interpret the biological implications of miRNA targets, Gene Ontology (GO) enrichment analysis was conducted via the clusterProfiler package. Analyses spanned three GO domains:

- **Biological Processes (BP)**: Identified pathways and processes potentially modulated by miRNA targets.
- **Molecular Functions (MF)**: Determines the biochemical activities associated with target genes.
- **Cellular Components (CC)**: Located the subcellular localizations pertinent to miRNA target functions.

Enrichment significance was ascertained using the Benjamini-Hochberg method, with a q-value cutoff of 0.05.

##### Visualizing gene ontology results

High-resolution visualizations were generated to effectively communicate findings:

- **Dot Plots**: Displayed the top enriched GO terms across BP, MF, and CC categories.
- **Heatmaps and Volcano Plots**: Provided comprehensive views of expression patterns and significance landscapes.

All plots were saved in publication-quality formats (300 dpi) to ensure clarity and reproducibility.

## RESULTS

### Metadata Filtering

Out of 585 age samples including different age groups. There were 491 samples after omitting the not available values “NAs”. From the available data, we selected 269 Elderly and 101 Senile samples. These samples when merged with the micro RNA expression data based on submitter ids of samples in both files. Only 204 Elderly and 76 senile samples were left.

**Fig 4.1:**
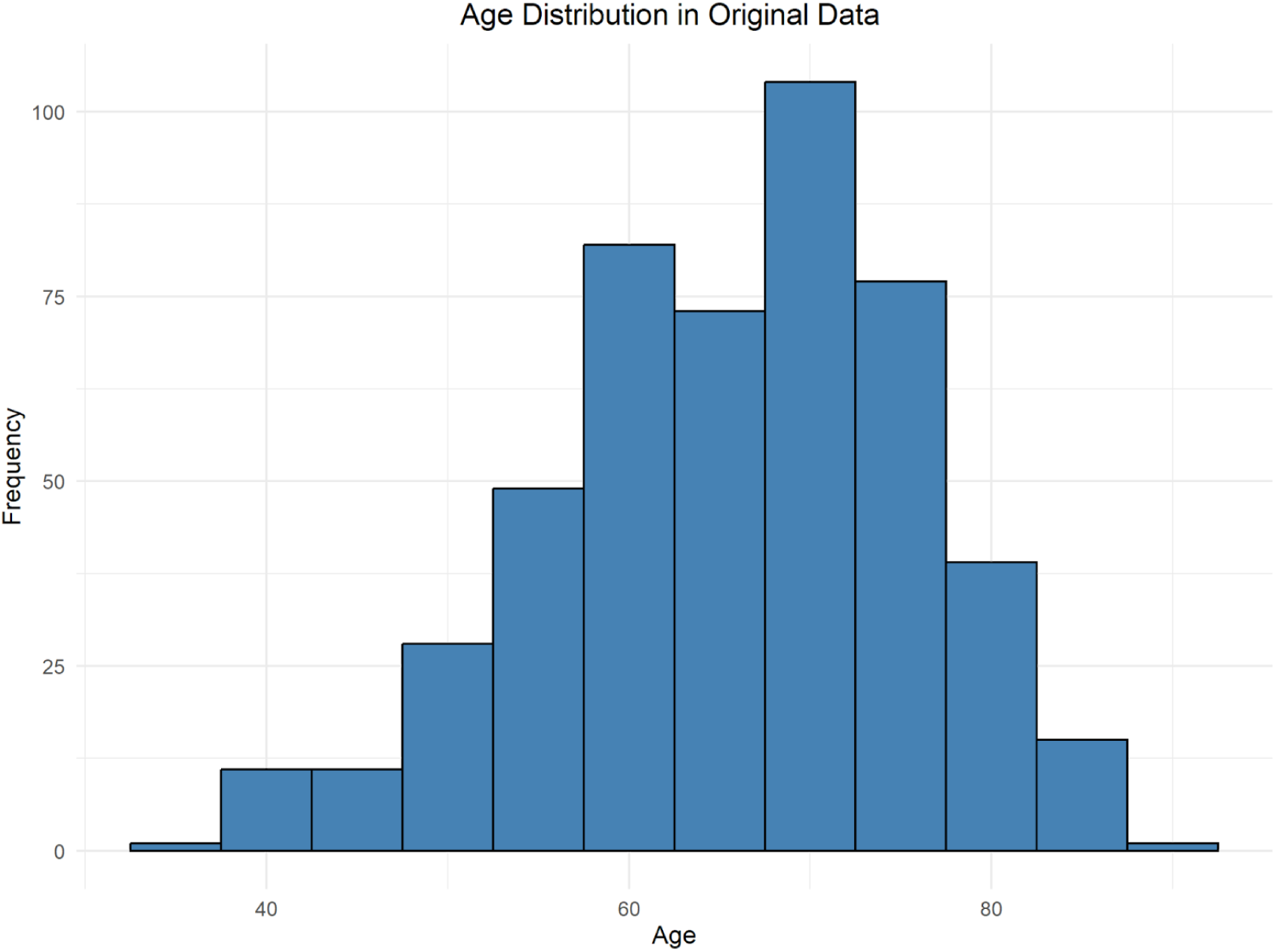
Age Distribution in raw data

**Fig 4.2:**
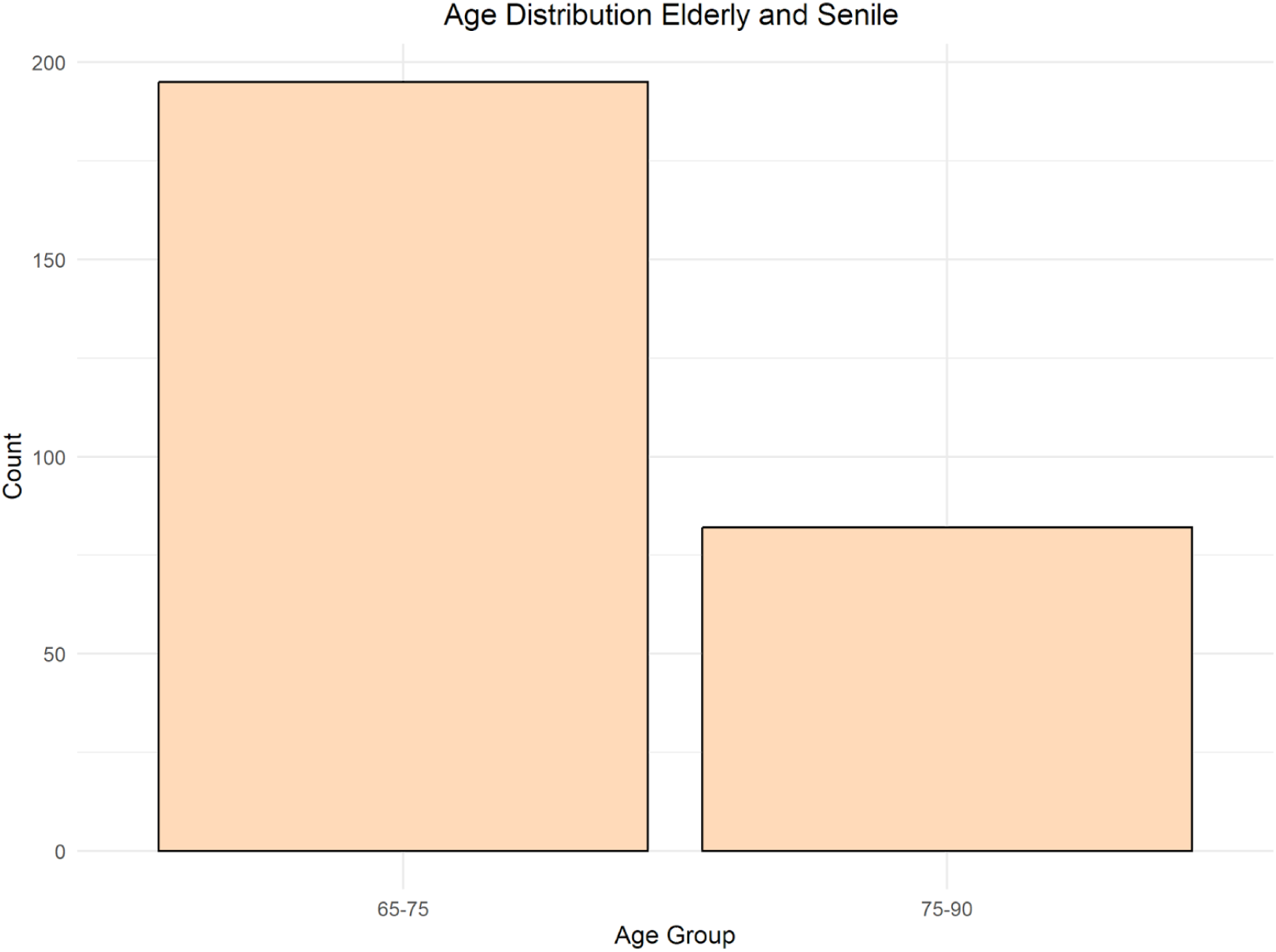
No of People of Elderly and Senile age group.

**Fig 4.3:**
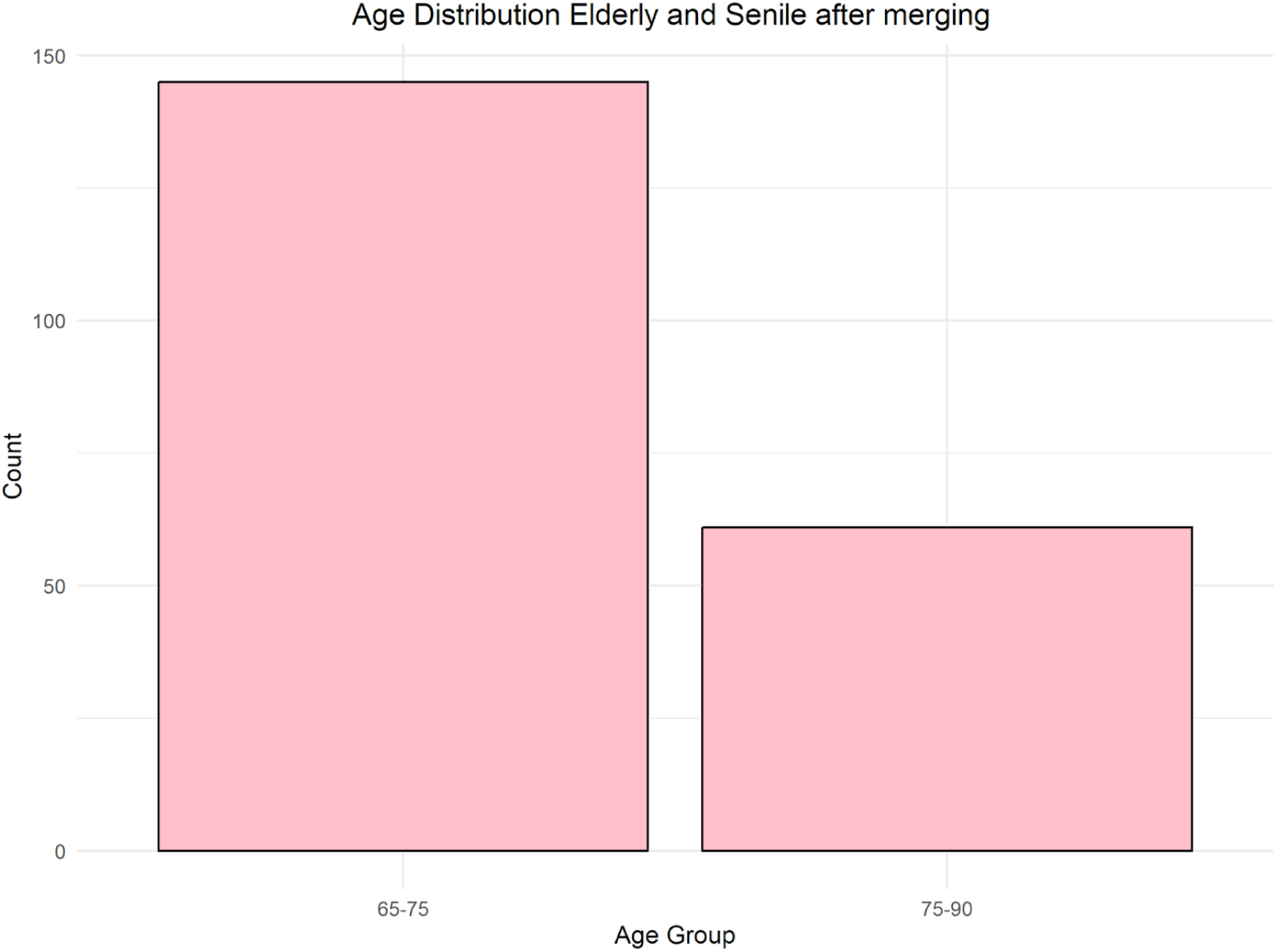
No of Elderly and Senile Samples left after Filtering and merging

### Differential Expression

801 differentially expressed miRNAs were identified using deseq2 out of 1881 miRNA expression profiles corresponding with 280 age samples.

To Further filter most significant miRNAs, the results of differential expression were filtered based on p-adjusted value (padj) of less than 0.05 and log2 fold change value greater than 1. As a result, 25 significant miRNAs were found.Only miRNA; hsa-mir-1911 was upregulated. Other 24 were down regulated.

List of Significant UP and Downregulated miRNAs, after applying based on padj value and log2foldchange(padj <0.05 and log2fold change > 1).

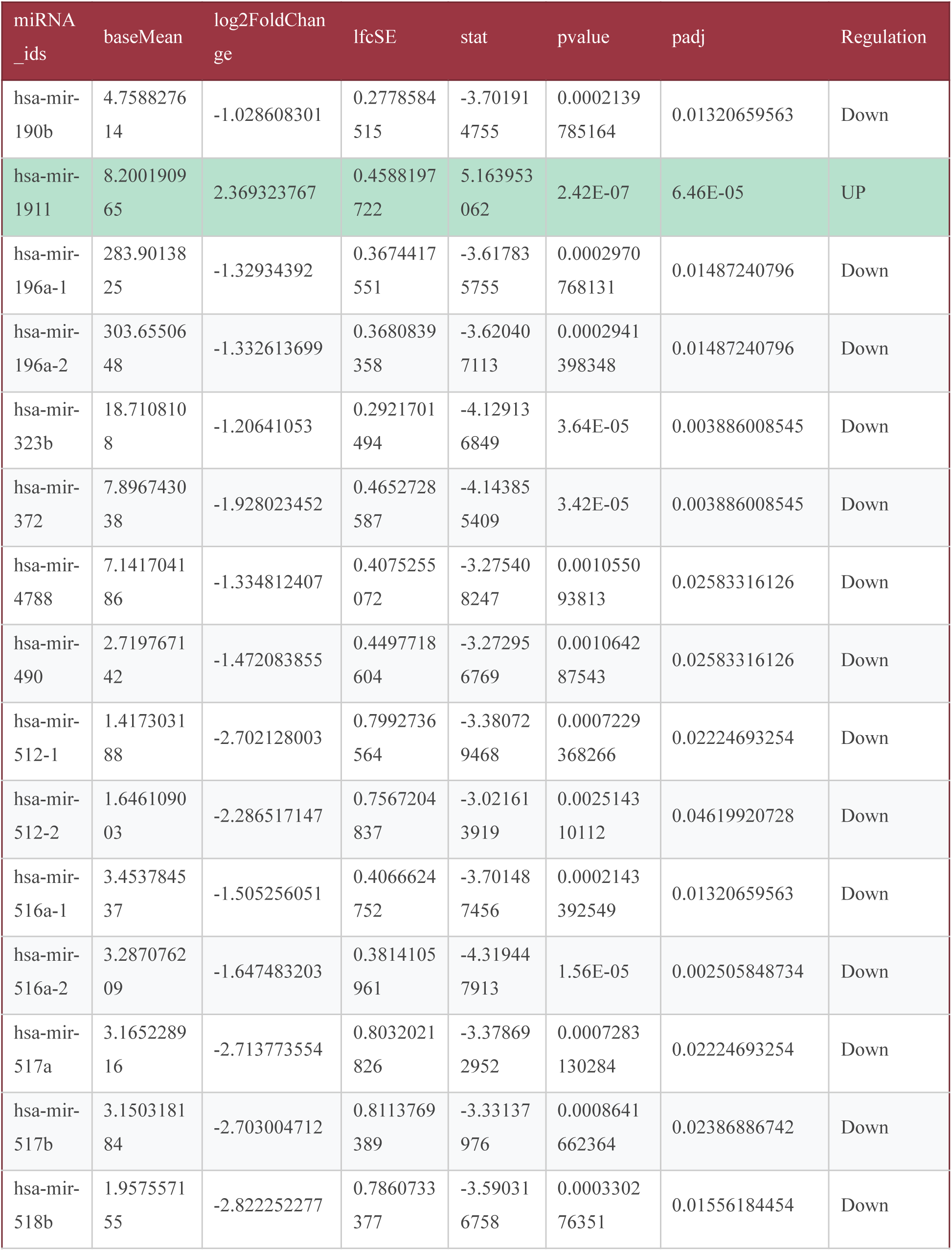
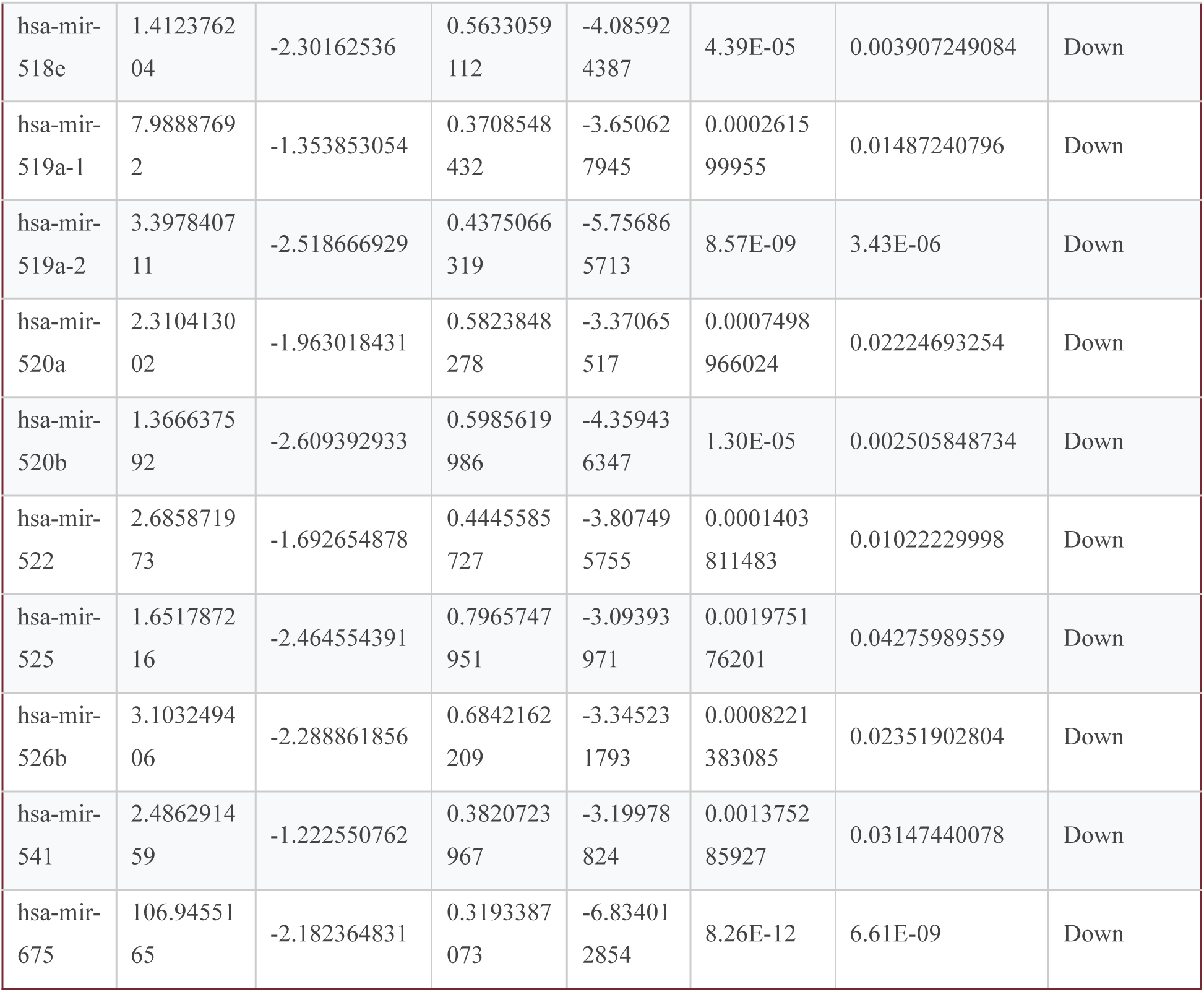

**Fig 4.4:**
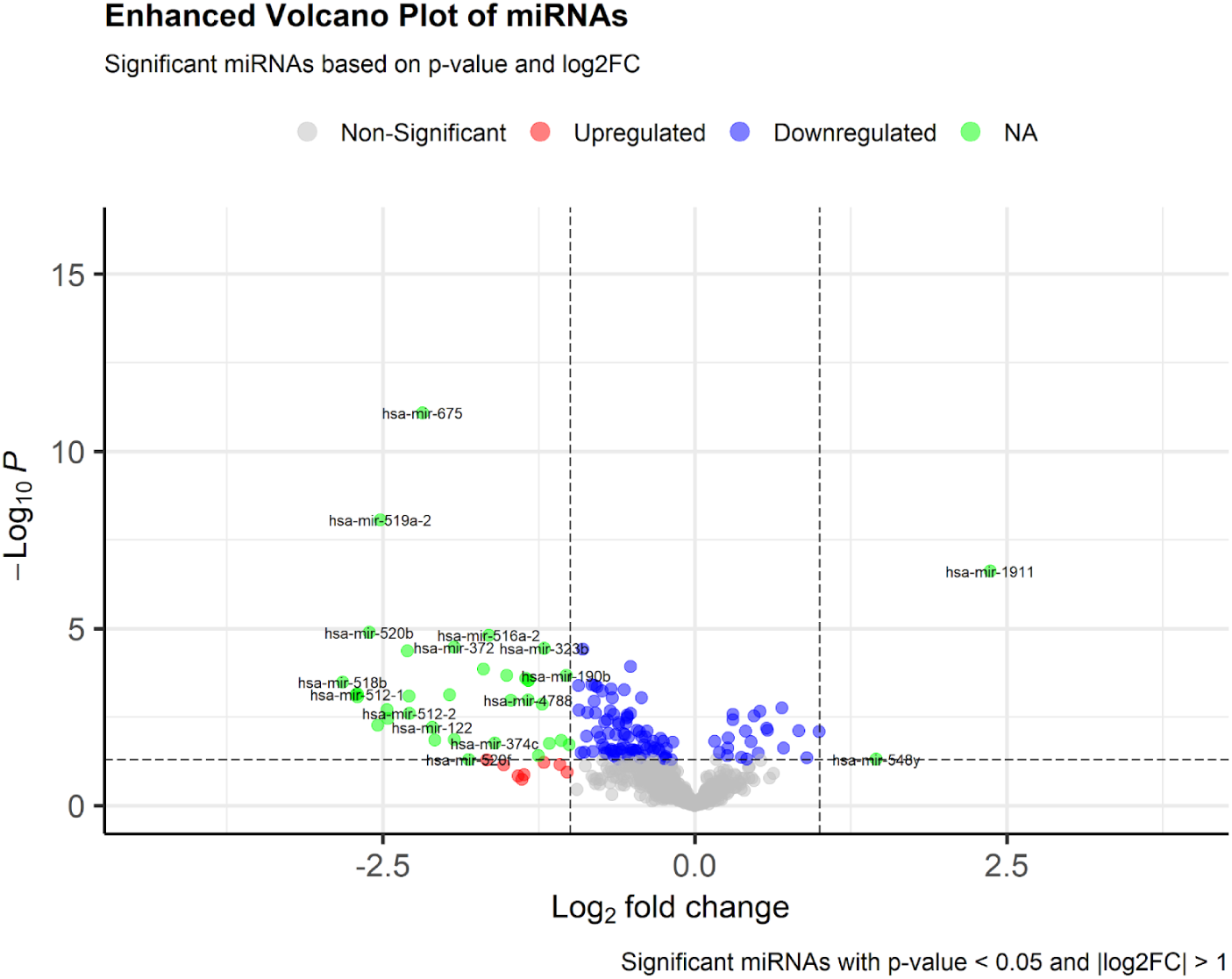
Volcano plot of miRNAs identified after DEA Via dseq2

list of significant miRNA With their related genes.

| mirbase_id | ensembl_gene_id | external_gene_name | description |
| --- | --- | --- | --- |
| hsa-mir-190b | ENSG00000215938 | MIR190B | microRNA 190b [Source:HGNC Symbol;Acc:HGNC:33656] |
| hsa-mir-1911 | ENSG00000222715 | MIR1911 | microRNA 1911 [Source:HGNC Symbol;Acc:HGNC:35395] |
| hsa-mir-196a-1 | ENSG00000210741 | MIR196A1 | microRNA 196a-1 [Source:HGNC Symbol;Acc:HGNC:31567] |
| hsa-mir-196a-2 | ENSG00000207924 | MIR196A2 | microRNA 196a-2 [Source:HGNC Symbol;Acc:HGNC:31568] |
| hsa-mir-323b | ENSG00000208004 | MIR323B | microRNA 323b [Source:HGNC Symbol;Acc:HGNC:38349] |
| hsa-mir-372 | ENSG00000199095 | MIR372 | microRNA 372 [Source:HGNC Symbol;Acc:HGNC:31786] |
| hsa-mir-490 | ENSG00000207597 | MIR490 | microRNA 490 [Source:HGNC Symbol;Acc:HGNC:32075] |
| hsa-mir-512-1 | ENSG00000207645 | MIR512-1 | microRNA 512-1 [Source:HGNC Symbol;Acc:HGNC:32090] |
| hsa-mir-512-2 | ENSG00000207644 | MIR512-2 | microRNA 512-2 [Source:HGNC Symbol;Acc:HGNC:32091] |
| hsa-mir-516a-1 | ENSG00000207767 | MIR516A1 | microRNA 516a-1 [Source:HGNC Symbol;Acc:HGNC:32130] |
| hsa-mir-516a-2 | ENSG00000207620 | MIR516A2 | microRNA 516a-2 [Source:HGNC Symbol;Acc:HGNC:32131] |
| hsa-mir-517a | ENSG00000207734 | MIR517A | microRNA 517a [Source:HGNC Symbol;Acc:HGNC:32111] |
| hsa-mir-517b | ENSG00000207837 | MIR517B | microRNA 517b [Source:HGNC Symbol;Acc:HGNC:32115] |
| hsa-mir-518b | ENSG00000207862 | MIR518B | microRNA 518b [Source:HGNC Symbol;Acc:HGNC:32106] |
| hsa-mir-5<br>18e | ENSG000002<br>07987 | MIR518E | microRNA 518e [Source:HGNC Symbol;Acc:HGNC:32119] |
| hsa-mir-5<br>19a-1 | ENSG000002<br>07992 | MIR519A1 | microRNA 519a-1 [Source:HGNC Symbol;Acc:HGNC:32128] |
| hsa-mir-5<br>19a-2 | ENSG000002<br>84362 | MIR519A2 | microRNA 519a-2 [Source:HGNC Symbol;Acc:HGNC:32132] |
| hsa-mir-5<br>20a | ENSG000002<br>07594 | MIR520A | microRNA 520a [Source:HGNC Symbol;Acc:HGNC:32099] |
| hsa-mir-5<br>20b | ENSG000002<br>07722 | MIR520B | microRNA 520b [Source:HGNC Symbol;Acc:HGNC:32105] |
| hsa-mir-5<br>22 | ENSG000002<br>83685 | MIR522 | microRNA 522 [Source:HGNC Symbol;Acc:HGNC:32127] |
| hsa-mir-5<br>25 | ENSG000002<br>07711 | MIR525 | microRNA 525 [Source:HGNC Symbol;Acc:HGNC:32102] |
| hsa-mir-5<br>26b | ENSG000002<br>07580 | MIR526B | microRNA 526b [Source:HGNC Symbol;Acc:HGNC:32100] |
| hsa-mir-5<br>41 | ENSG000002<br>16179 | MIR541 | microRNA 541 [Source:HGNC Symbol;Acc:HGNC:33650] |
| hsa-mir-6<br>75 | ENSG000002<br>88367 | MIR675 | microRNA 675 [Source:HGNC Symbol;Acc:HGNC:33351] |
| hsa-mir-6<br>75 | ENSG000002<br>84010 | MIR675 | microRNA 675 [Source:HGNC Symbol;Acc:HGNC:33351] |

**Fig 4.5:**
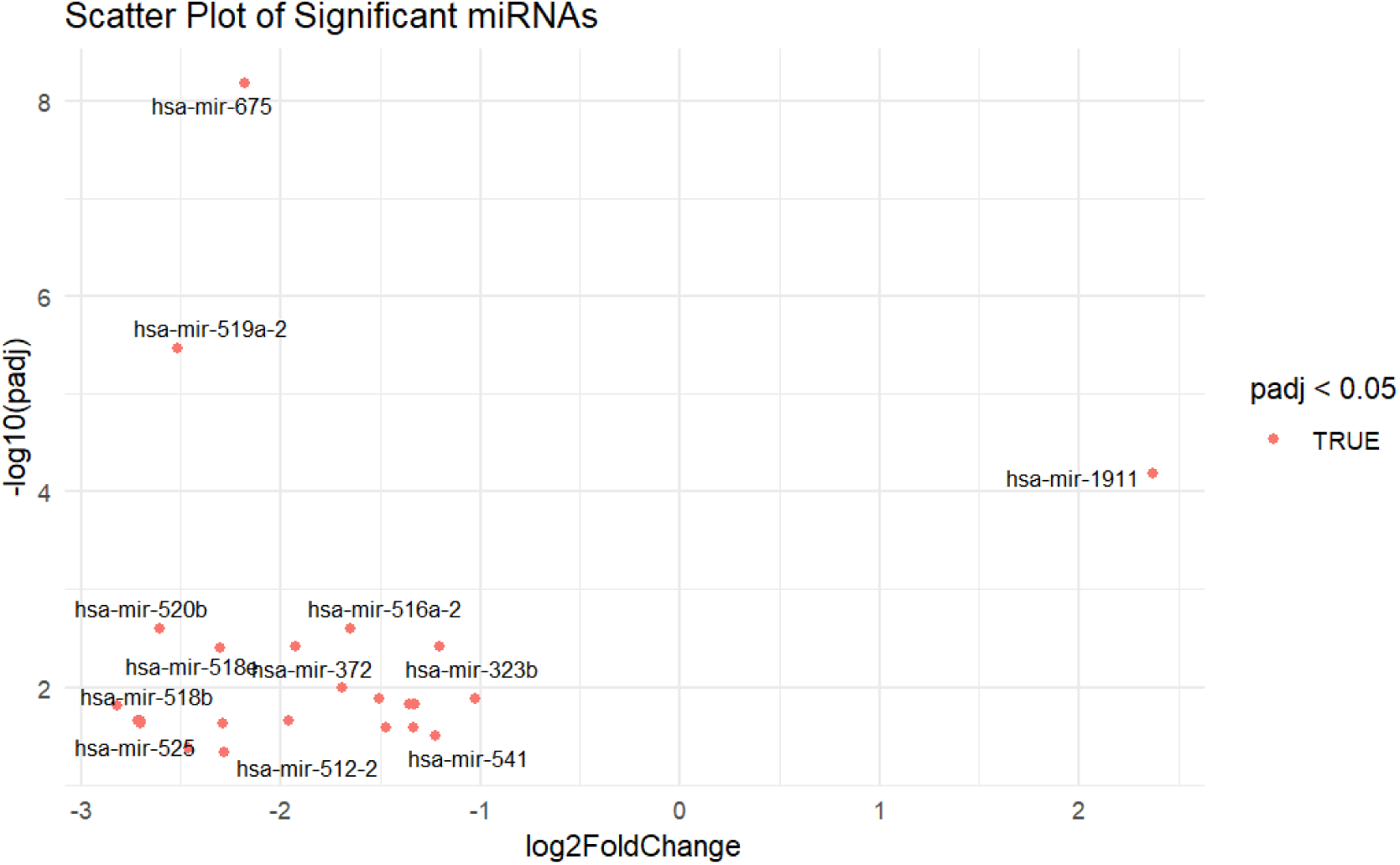
Scatter plot showing significant miRNAs identified

#### ML_Results

Important miRNAs found by random forest, hsa-mir-181b.1, hsa.mir.542, hsa.mir.450b, hsa.mir.584, hsa.mir.21.

**Fig 4.6:**
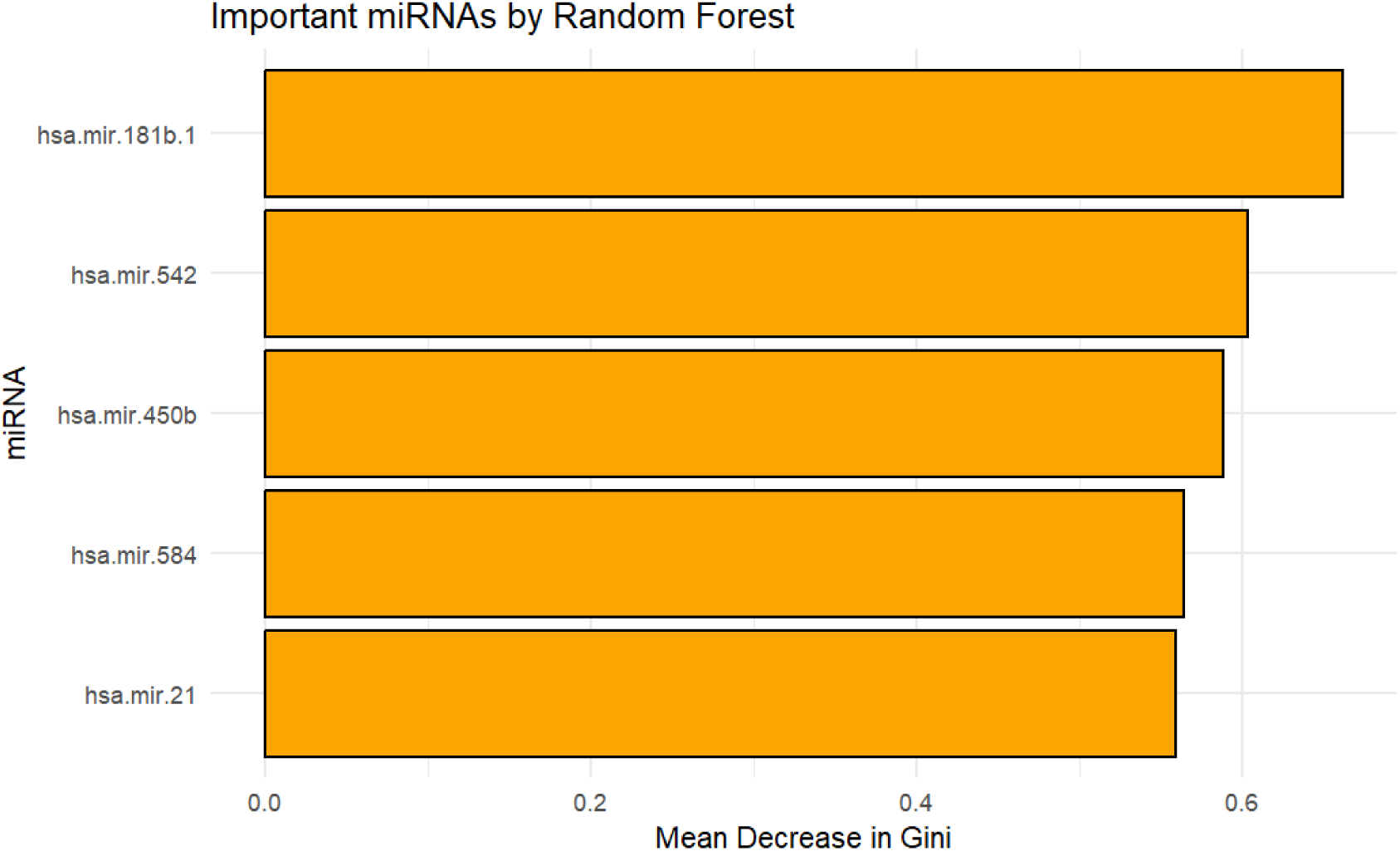
Important miRNAs via mean decrease score

**Fig 4.7:**
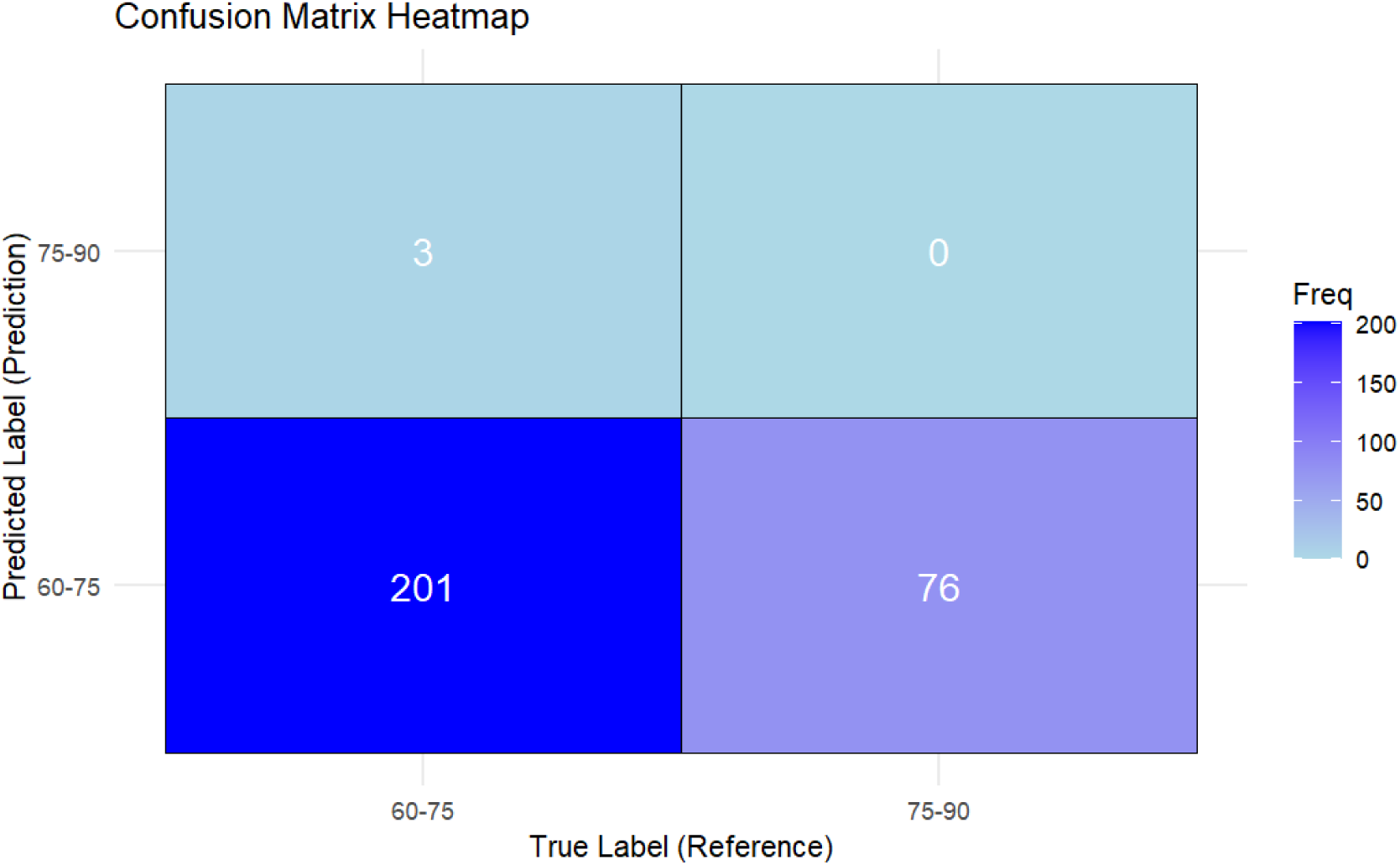
Confusion matrix of Random forest model

### Functional Enrichment

#### Biological_Process

Biological Processes identified “miRNA-mediated gene silencing by destabilizing mRNA”.This biological process was significant with the padj value of 0.033 and is associated with 3 genes.

**Fig 4.8:**
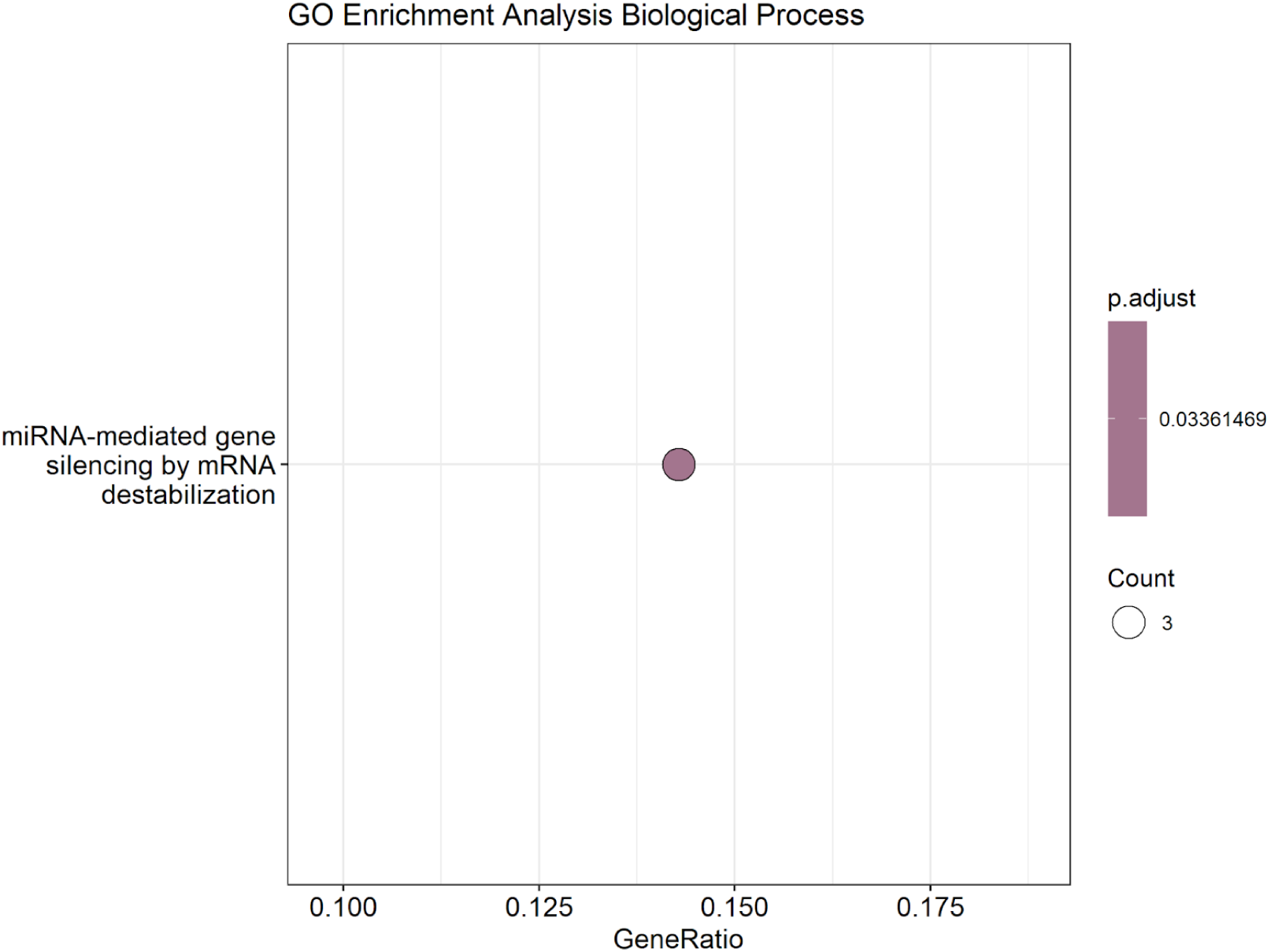
GO enrichment biological Process showing Biological processes related to 3 most related genes

#### Molecular_Function

Molecular functions of mRNA base pairing translational repressor activity and translation repressor activity showed lower padj value and more number of genes while translation regulator activity was associated with less number of genes and higher padj value.

**Fig 4.9:**
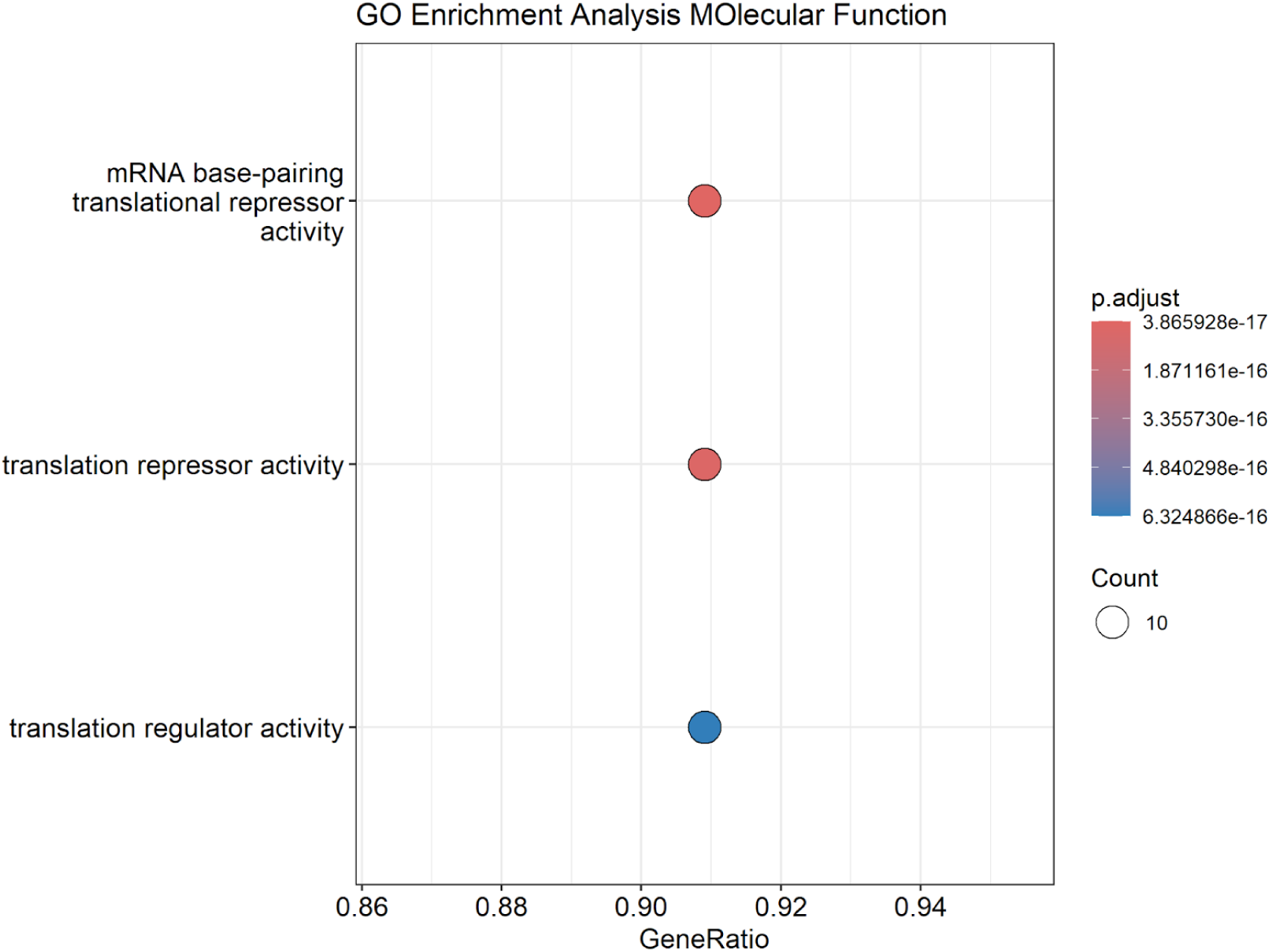
GO enrichment showing molecular functions related of miRNA associated genes

#### Cellular_Component

Functional Enrichment showed two cellular component complexes RNAi effector and RISC Complex associated with 7 genes but with a high padj value of 5.36 e-11.

**Fig 5.0:**
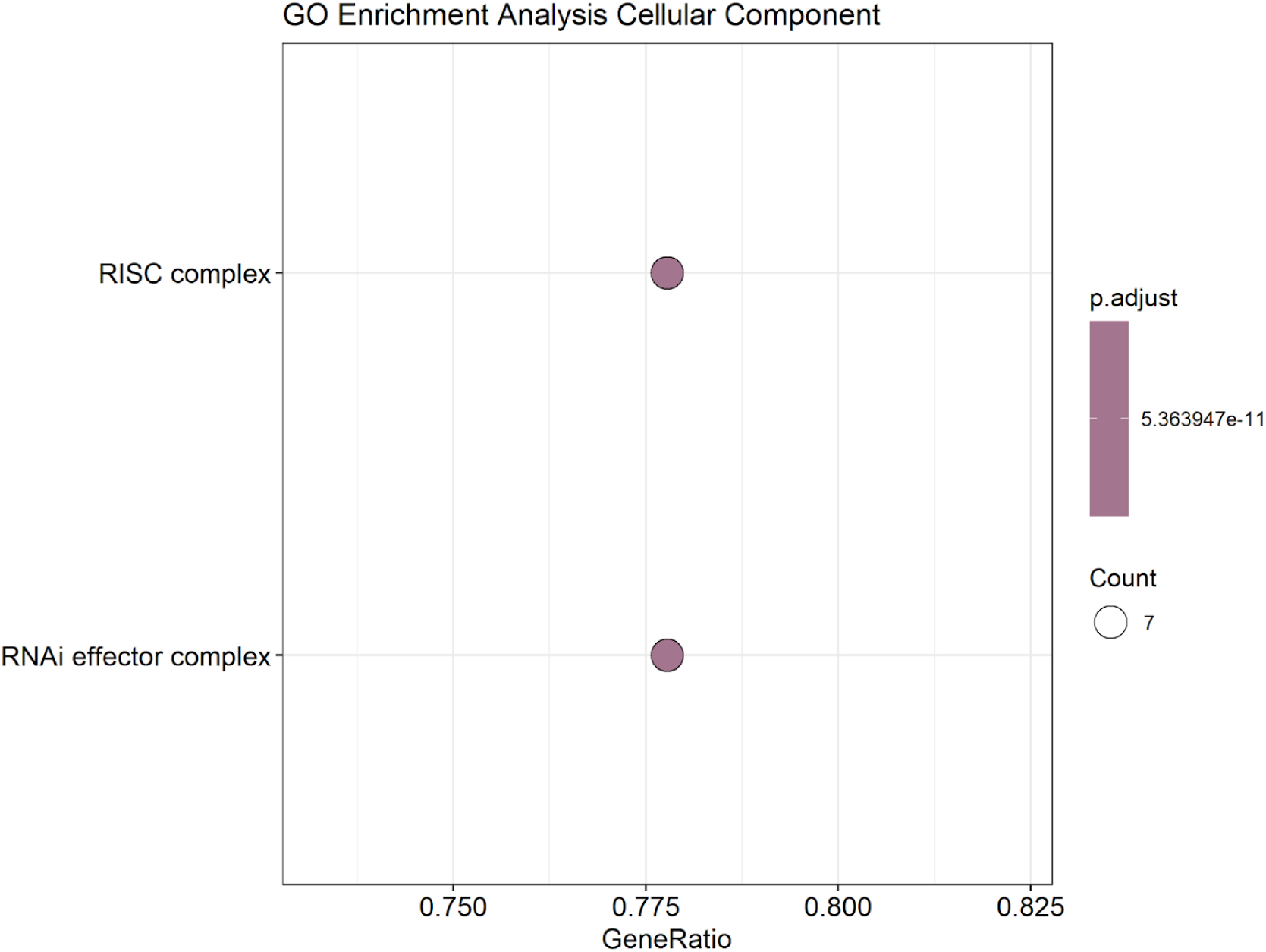
Cellular Components related to identified significant micro RNAs

## DISCUSSION

The identification of age-associated miRNAs offers valuable insights into the molecular mechanisms underlying age-related biological processes. In this study, a rigorous filtering of metadata and miRNA expression data led to the inclusion of 280 samples across two distinct age groups: elderly and senile. This meticulous data processing ensured the reliability of the downstream analyses.

Differential expression analysis revealed 801 miRNAs as differentially expressed out of 1,881 miRNA profiles, with further stringent filtering highlighting 25 significant miRNAs (padj < 0.05, log2 fold change > 1). Notably, hsa-mir-1911 was the only upregulated miRNA, while the remaining 24 were significantly downregulated. These findings align with existing literature, suggesting that miRNAs play a critical role in age-related cellular regulation, including apoptosis, senescence, and inflammation. For example, downregulated miRNAs like hsa-mir-196a and hsa-mir-323b have been previously implicated in processes such as stem cell differentiation and immune modulation.

The unique upregulation of hsa-mir-1911 warrants special attention. Previous studies suggest its involvement in regulatory pathways related to aging, potentially linked to cellular stress response and mitochondrial function. This miRNA could serve as a candidate biomarker for age-related pathologies, particularly in distinguishing aging phenotypes between elderly and senile individuals.

Interestingly, the observed downregulation of miRNAs such as hsa-mir-675, hsa-mir-372, and hsa-mir-519a points to their potential roles in suppressing tumor-suppressive or pro-apoptotic pathways during aging. These miRNAs, along with their identified target genes, may serve as key regulatory elements in aging-related conditions such as neurodegeneration and age-associated cancers.

Additionally, the overlap of miRNA targets with known pathways highlights the interconnected nature of aging biology, with several targets involved in processes such as inflammation, oxidative stress, and metabolic regulation. Future studies should focus on validating these findings in larger, independent datasets and elucidating the functional roles of these miRNAs through experimental approaches.

The limited number of upregulated miRNAs compared to the downregulated ones emphasizes the need for further investigation into mechanisms that drive the suppression of miRNA expression during aging. This phenomenon might reflect age-related epigenetic modifications or transcriptional repression.

### 5.1 Overview of miRNAs Identified in This Study

#### miR-181b

Tumor Suppressor RoleThis research evaluates the tumor-limiting characteristics of miR-181b within non-small cell lung cancer (NSCLC). The researchers reported that NSCLC tissues contained significantly less miR-181b in comparison to normal lung tissues during the study. Low miR-181b expression correlated with extensive tumors and multiple lymph node metastases in addition to advanced disease classification that suggested the protein’s impact on aggressive patient outcomes. Furthermore survival analysis showed that patients with decreased miR-181b expression exhibited worse survival rates both for complete survival and disease-free state thus qualifying it as a predictive marker for NSCLC. Independent analysis of both univariate and multivariate results demonstrated that miR-181b expression levels function as an independent indicator of unfavorable disease prognosis. The results indicate that NSCLC tumor progression depends on reduced miR-181b expression levels which provide potential value for NSCLC outcome prediction. (Yang et al., 2013)

#### Hsa-miR-542-3p

MiR-542-3p functions as a tumor suppressor in NSCLC. miR-542-3p achieves its suppressive activity through 3′UTR targeting of FTSJ2 mRNA leading to FTSJ2 mRNA and protein level elevation. The results demonstrate that miR-542-3p functions as a tumor suppressor which inhibits NSCLC cancer progression in NSCLC (Liu et al., 2017).

#### Hsa-mir-21

Research identifies NSCLC oncogene MiR-21 as overexpressed in multiple cancers where it contributes to proliferation along with apoptosis suppression and invasion and metastatic behaviors. The overexpression pattern exists in different malignancies because it supports cell proliferation while inhibiting apoptosis and enables invasion and metastasis. Studies have demonstrated the connection between elevated miR-21 levels and unluckier clinical outcomes in NSCLC patients together with proven anti-NSCLC effects when miR-21 is blocked. The investigation reveals PTEN as a target molecule of miR-21 that likely leads to decreased PTEN protein levels to contribute to tumor progression (Zhang et al., 2010).

#### Hsa-mir-450b

The research shows MiR-450b serves as an inhibitor of cancer cell growth in NSCLC. Consequences of reduced MiR-450b-5p expression suggest a tumor-suppressing mechanism because lung adenocarcinoma cells experience decreased malignant potential. The clinical value of MiR-450b-5p as a diagnostic tool and prognostic indicator needs additional studies to understand its precise role in NSCLC diagnostics.(Wu et al.,2022)

#### Hsa-mir-584

miR-584 functions as a tumor suppressor in non-small cell lung cancer (NSCLC). Tumor tissue samples show significant downregulation of miR-584-5p expression which associates with more advanced stages as well as lymph node metastasis. This microRNA directly targets MMP-14 to inhibit cell migration and invasion in cancer cells. The expression of MMP-4 and Slug proteins that facilitate metastasis and EMT is also reduced by its effect. Lung cancer progression suppression by miR-584-5p creates strong support for its implementation as both a biomarker and therapeutic target in NSCLC research Guo et al. (2019).

## CONCLUSION

This study successfully identified 25 significant age-associated miRNAs through differential expression analysis of 280 samples. The results emphasize the predominance of downregulated miRNAs, with hsa-mir-1911 as the sole upregulated candidate, highlighting its potential as a biomarker for aging. The findings provide a foundation for understanding the miRNA-mediated regulatory mechanisms involved in aging and pave the way for future research to explore therapeutic interventions targeting miRNA pathways in age-related diseases.

In conclusion, miRNAs represent promising biomarkers and therapeutic targets in the context of aging. By integrating these results with functional studies and pathway analyses, future research can unlock novel strategies to combat age-associated disorders and improve the quality of life in aging population.

